# Differential role for VPS4 isoforms in cytokinetic abscission confers a regulatory function for monomeric VPS4A and VTA1 in mammalian cells

**DOI:** 10.1101/2023.09.11.557113

**Authors:** Inbar Dvilansky, Yarin Atlaras, Dikla Nachmias, Natalie Elia

## Abstract

Mutations in the human AAA-ATPase VPS4 isoform, VPS4A, cause severe Neurodevelopmental defects and Congenital Dyserythropoietic Anemia (CDA). VPS4 is a crucial component of the ESCRT system, which drives membrane remodeling in numerous cellular processes, including receptor degradation, cell division, and neural pruning. Notably, while most organisms encode for a single VPS4 gene, human cells have two VPS4 paralogs, namely VPS4A and VPS4B, but the functional differences between these paralogs is mostly unknown. Here, we set out to investigate the role of the human VPS4 paralogs in cytokinetic abscission using a series of knockout cell lines. We found that VPS4A and VPS4B hold both overlapping and distinct roles in abscission. VPS4A depletion resulted in a severe abscission delay, which was fully rescued by VPS4A expression but only partially rescued by VPS4B overexpression. Unexpectedly, expressing a monomeric-locked VPS4A mutant also partially rescued the abscission delay in VPS4A KO cells and bound the abscission checkpoint proteins CHMP4C and ANCHR. Depletion of VTA1, a co-factor of VPS4, disrupted VPS4A- ANCHR interactions and accelerated abscission, indicating a role for VTA1 in the abscission checkpoint. Our findings reveal a dual role for VPS4A in abscission, one that is canonical and can be compensated by VPS4B, and another that is regulatory and is mediated by its monomeric form. These observations provide a potential mechanistic explanation for the neurodevelopmental defects and other related disorders reported in VPS4A-mutated patients with a fully functional VPS4B paralog.

## INTRODUCTION

VPS4 is one of the core components of the Endosomal Sorting Complex Required for Transport (ESCRT) membrane remodeling system, which is currently recognized as one of the most basic cellular machineries for driving membrane fission in cells. As such, VPS4 is essential for in vitro reconstitution of ESCRT-induced membrane fission and is involved in all canonical ESCRT mediated processes in eukaryotes, including in multivesicular body (MVB) formation, release of retroviruses from the cell surface, nuclear envelope sealing, plasma membrane repair, neural pruning, and scission of daughter cells during the last stages of cytokinesis (*1-3*). Additional non-canonical roles for VPS4 in centrosomes have also been reported (*4, 5*). Finally, mutations in VPS4 genes were associated with different neuro-pathologies and cancer (*6-9*). Therefore, understanding the basis for VPS4 function in cells is of great interest.

VPS4 is an AAA-ATPase that assembles into functional hexamers or dodecamers that hydrolyze ATP (*10, 11*). This functional hexamer is stabilized by binding to its co-factor VTA1, which dimerizes to form a bridge that connects two adjacent VPS4 subunits (*11, 12*). While yeast cells encode for a single VPS4 protein, mammalian cells encode for two VPS4 homologs, namely VPS4A and VPS4B. Given the high sequence similarity of these paralogs (81% amino acids identity), they were traditionally thought to have redundant functions in cells (*8, 13*). However, recent evidence suggests that the cellular functions of VPS4A and VPS4B do not fully overlap. First, mutations in VPS4A in patients carrying normal VPS4B caused severe diseases associated with structural brain abnormalities, neurodevelopmental defects, cataract, growth defects, and Congenital Dyserythropoietic Anemia (CDA) (*6, 7*). Second, while VPS4A was reported to function as a tumor suppressor, VPS4B exhibited pro-or anti-oncogenic activities in different cancers (*8, 9*). Nevertheless, depleting VPS4A in cancer cells with loss of VPS4B led to synthetic cell death, suggesting at least partial overlap between these isoforms (*8*). Third, VPSB was previously shown to be more crucial for HIV viral release than VPS4A, and we recently substantiated these findings and showed that depletion of VPS4A or VPS4B has differential effects on HIV-viral release, suggesting a unique property for each isoform (*14, 15*). Collectively, these reports raise possibility that the two VPS4 isoforms hold overlapping and unique functions in cells stressing the need to investigate the functional differences of these two isoforms in cells.

Cytokinetic abscission provides an excellent platform for dissecting the role of VPS4 paralogs. During abscission, components of the ESCRT-III complex assemble into helical filaments at the intercellular bridge that connect the two dividing daughter cells, and drive constriction and fission of the membranes connecting these cells to complete the division process. VPS4 participates in several crucial aspects of this process. First, it is involved in the exchange of ESCRT-III monomers within the filament during filament assembly (*16*). Second, it plays a role in ESCRT-III depolymerization and membrane fission that terminates abscission (*17, 18*). Third, it interacts with proteins of the AuroraB abscission checkpoint (ANCHR and CHMP4C), which regulates abscission timing (*19, 20*). Notably, both paralogs localized to the intercellular bridge when overexpressed in cells (*16, 18*), suggesting their functional involvement in abscission. Therefore, VPS4 exhibits multiple functions in abscission, with both VPS4 paralogs potentially contributing to these functions.

Here we set out to investigate the functions of the two VPS4 isoforms encoded in mammalian cells during cytokinetic abscission by knocking out specific VPS4 components. We found that although cells express higher levels of VPS4B compared to VPS4A, depletion of VPS4A leads to a considerably more severe abscission delay phenotype than depletion of VPS4B. Stochastic Optical Reconstruction Microscopy (STORM) measurements of the density of ESCRT-III proteins at the intracellular bridge pointed to the involvement of VPS4A and VPS4B at different abscission stages, supporting a differential role for VPS4 isoforms. Interestingly, a VPS4A mutant defected in hexamerization (mVPS4A) expressed in VPS4A KO cells, arrived at the cytokinetic bridge, and partially rescued the abscission delay phenotype. Unexpectedly, depleting the VPS4 co-factor VTA1, accelerated abscission. In WT cells, VTA1 was found to interact with both CHMP4C and ANCHR, suggesting its role in the abscission checkpoint. mVPS4A was also found to interact with CHMP4C and ANCHR, but its interaction with ANCHR was abolished in VTA1 KO cells, suggesting the formation of a VPS4A-VTA1-ANCHR complex at the abscission checkpoint. Collectively, our data indicate that VPS4 A and B isoforms have distinct functions remodeling the ESCRT-III filament during mediated cytokinetic abscission, that could be partly compensated by the presence of one isoform. However, the regulatory role of VPS4 in abscission is mediated by the VPS4A isoform in its monomeric form and is dependent on VTA1 binding. Notably, the regulatory role of VPS4A shown here provide for the first time a mechanistic explanation for the previously reported direct involvement of VPS4A (but not VPS4B) in disease development (*6, 7*).

## RESULTS

To dissect the role of VPS4 isoforms in cytokinesis, we monitored abscission in VPSA and VPS4B KO cells recently generated in our laboratory (Fig. S1A) (*14*). Attempts to generate cells depleted of both VPS4A and VPS4B resulted in cell death, suggesting that at least one VPS4 isoform is needed for cell viability, as previously suggested (*8*). Depleting either VPS4A or VPS4B caused a delay in abscission and an increase in the percentage of cells that failed to complete abscission but did not affect the formation or morphology of the intercellular bridge (Fig. 1A). Notably, VPS4A KO cells exhibited a considerably more severe abscission delay, which could be fully rescued by exogenous VPS4A expression (Fig. 1A, B). Exogenous VPS4B expressed in VPS4A KO cells arrived at the intercellular bridge and partially rescued the abscission delay (Fig. 1B, Fig. S1C), indicating that VPS4B cannot fully compensate for the loss of VPS4A in abscission. Measurements of the total levels of VPS4 paralogs in cells revealed that VPS4B is significantly more abundant than VPS4A (∼ 5:1, respectively) (Fig. S1B). These findings suggests that VPS4A is the more prominent paralog in abscission, despite its low cellular abundance.

**Figure 1:**
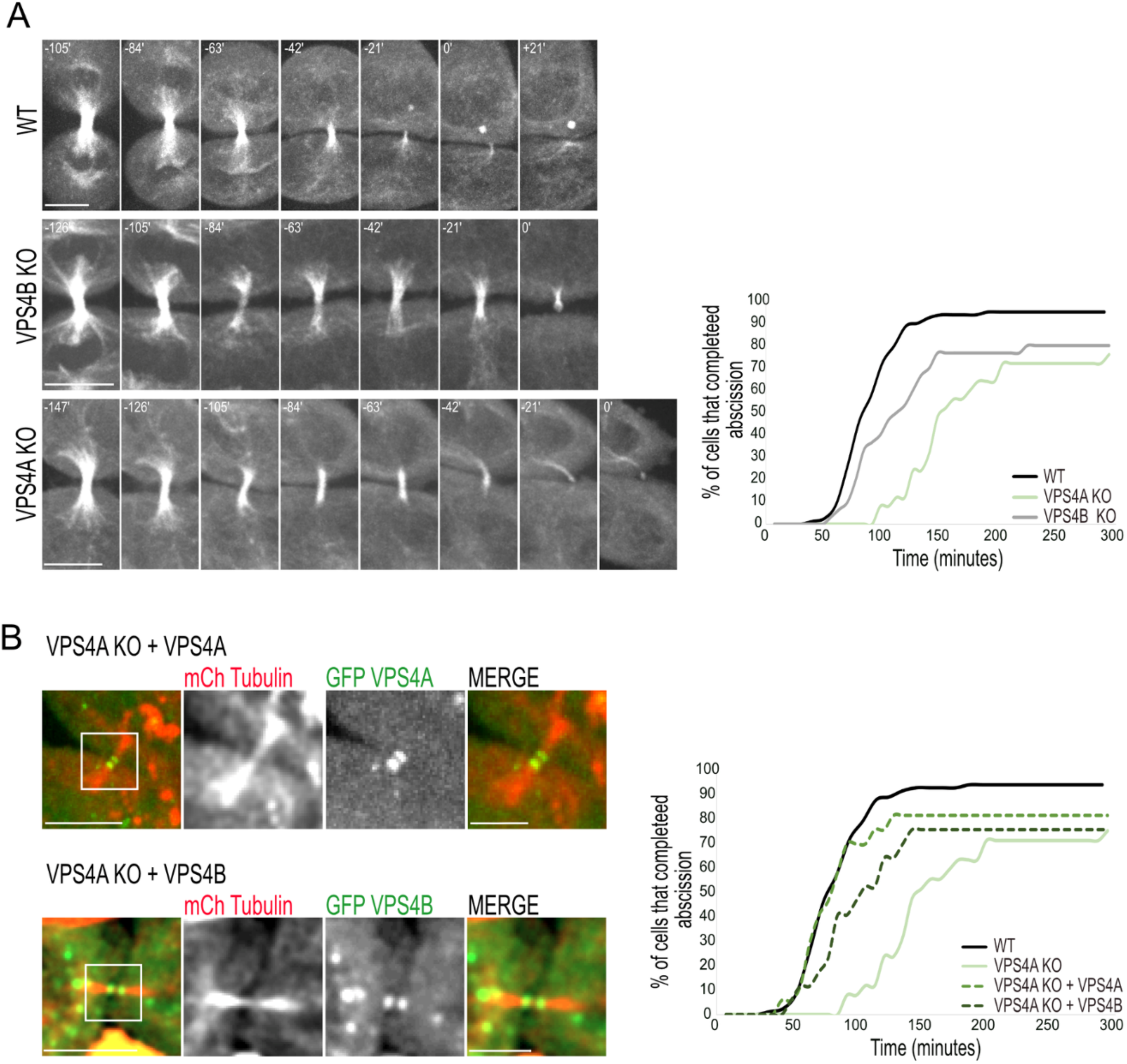
Knock-out of VPS4A causes a stronger abscission delay phenotype than VPS4B knock-out. (**A**) Live cell imaging of WT (top panel), VPS4B KO (middle panel), and VPS4A KO (bottom panel), transfected with GFP-tubulin. Z slices were captured at 7-minute intervals using a confocal spinning-disk microscope. Maximum-intensity projections of representative cells at various time points during cytokinesis are shown. The abscission time was set as time = 0 and was defined as the time of the first microtubule bridge cleavage event. A cumulative plot showing the duration of cytokinesis (from cleavage furrow formation to microtubule bridge resolution) of all cells recorded is shown on the right. Averaged duration time of abscission was longest in VPS4A KO cells compared to WT cells and VPS4B KO cells (VPS4A KO 155 ± 46 minutes, n = 25; VPS4B KO 106 ± 37 minutes, n = 30; WT 90 ± 25 minutes, n = 76). Scale, 10 µm. (**B**) The abscission delay observed in VPS4A KO cells is fully rescued upon overexpression of VPS4A and partially rescued upon overexpression of VPS4B. VPS4A KO cells were co-transfected with mCherry-Tubulin (red) together with either GFP-VPS4A (top panel, green) or GFP-VPS4B (bottom panel, green), and imaged 48 hours later, at 7-min intervals. Left panels: zoom-out images (scale, 10 µm). Zoom-in images of the white squares in left panels are shown on the right (scale, 5 µm). Shown are maximum-intensity projection images of representative cells. Note the arrival of VPS4A and VPS4B to the intercellular bridge in VPS4A KO cells. Complete movie series are provided in Sup. Movies 1 (top panel) and 2 (bottom panel). A cumulative plot showing the duration of cytokinesis (from cleavage furrow formation to microtubule bridge resolution) of all cells recorded is shown on the right (dashed lines) (VPS4A KO + GFP-VPS4A 85 ± 22 minutes, n = 17; VPS4A KO + GFP-VPS4B 100 ± 28 minutes, n = 15). WT and VPS4A KO cell data (reproduced from Fig. 1A) are shown for reference.

During cytokinetic abscission, components of the ESCRT-III complex assemble into high-ordered structures on the intracellular bridge (*18, 21, 22*). Structured Illumination Microscopy (SIM) imaging of the ESCRT-III proteins IST1 and CHMP4B in VPS4 KO cells revealed that the overall organization of the ESCRT-III filaments at the intercellular bridge is not perturbed upon depletion of either isoform, suggesting that one VPS4 isoform is sufficient for proper ESCRT-III organization (Fig. 2A, S1D). That said, quantification of IST1 density at the intercellular bridge using STORM analysis pointed to a differential role of VPS4 paralogs in abscission. In early stages, depletion of VPS4A, but not of VPS4B, caused accumulation of IST1 compared to WT cells, suggesting a role for VPS4A in early stages of abscission. In late stages, a decrease in IST1 densities was observed in WT cells, indicating either the removal of IST1 from the ESCRT-III filament or depolymerization of the ESCRT-III filament. While this phenotype was recapitulated in VPS4A KO cells, it was almost abolished in VPS4B KO cells (Fig. 2B). These results suggest that the VPS4A isoform is primarily involved in early stages of abscission, where it facilitates the exchange of ESCRT-III components, while the VPS4B isoform act in late stages, predominantly mediating depolymerization of ESCRT-III subunits. These data indicate that VPS4A and VPS4B have distinct, temporally separated functions in abscission.

**Figure 2:**
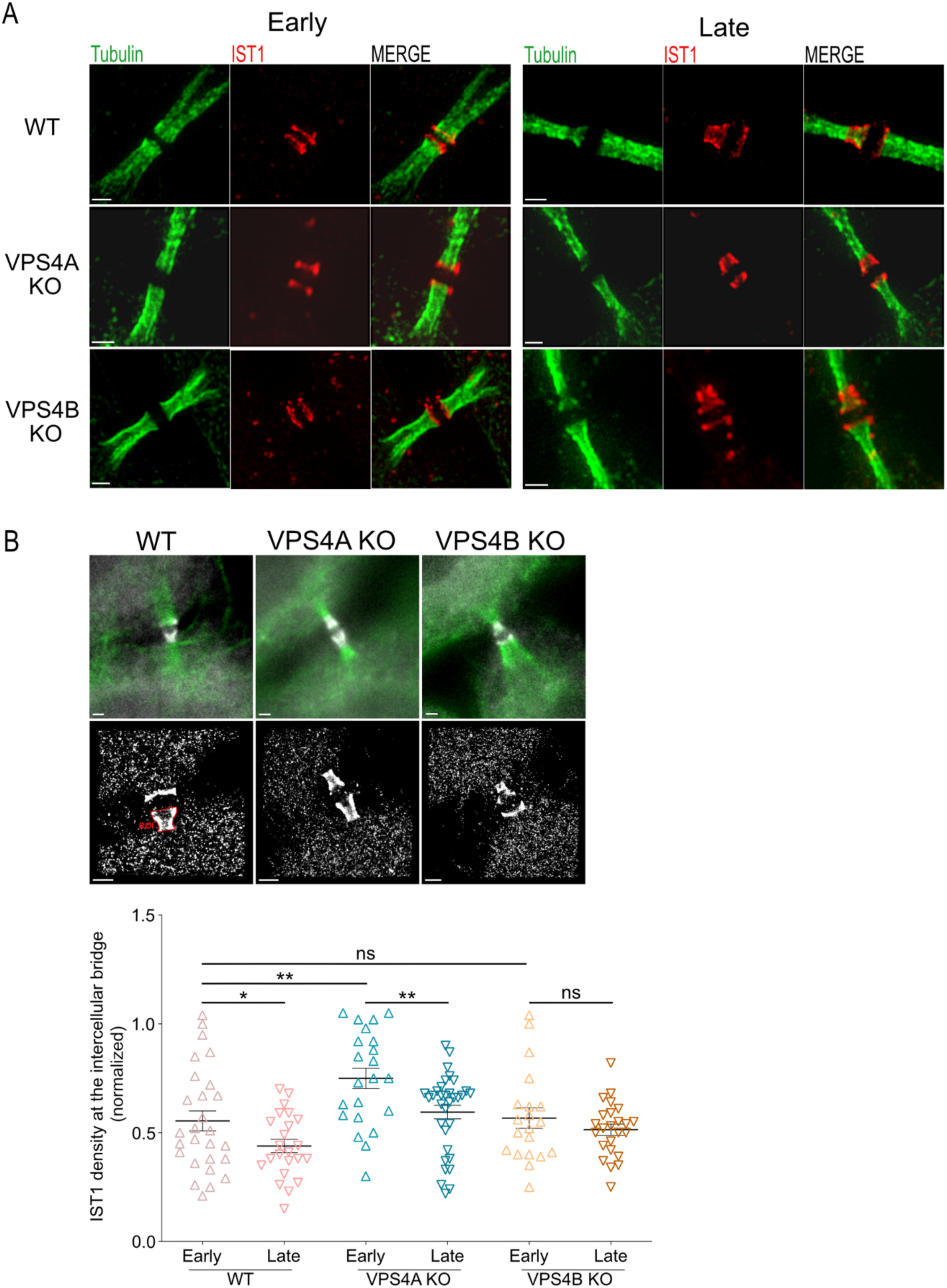
Differential effect of VPS4A and VPS4B depletion on the organization of ESCRT-III at the intercellular bridge. (**A**) SIM imaging shows normal ESCRT-III localization at the intercellular bridge in VPS4 KO cells. WT (top panel), VPS4A KO (middle panel), and VPS4B KO (bottom panel) cells were stained with anti-α-tubulin (green) and anti-IST1 (red) antibodies and imaged by SIM. Maximum projection images of early (left) and late (right) intercellular bridges are shown. Scale, 1µm. (**B**) STORM imaging shows different ESCRT-III densities at intercellular bridges of VPS4A and VPS4B KO cells. Cells were synchronized, stained with anti-α-tubulin and anti-IST1 antibodies and imaged by 2D STORM in epifluorescence mode (see methods section). Wide field images showing the intercellular bridge and IST1 staining (tubulin, green; IST1, white), and their respected STORM datasets of IST1 localizations are shown on top panels (scale, 1 µm). A scatter plot graph showing the density of IST1 at the intercellular of all cells measured. IST1 density was calculated by dividing the total number of localizations by the area of manually selected regions (ROIs) that include IST1 staining on one side of the bridge (red marking, left panel). Statistical analysis was performed for five independent experimental repeats, using two-tailed t-test (*p-value ≤ 0.05, **p-value ≤ 0.01. WT: early, n = 28, late, n = 24; VPSA KO: early, n = 22, late, n = 33; VPSB KO: early, n= 20, late, n=24). n refers to the number of ROIs.

In cells, VPS4 resides in its monomeric or dimeric forms in the cytosol, which assemble into active ATPase hexamers upon binding to membrane-bound ESCRT-III (*23*). To examine whether the abscission delay phenotype observed for VPS4A in abscission relies on hexamerization, we generated a VPS4A monomeric mutant based on a previously reported yeast VPS4 mutant that was found to be monomeric in biochemical assays (L145D, corresponds to L151D in yeast VPS4) (*24*). A fluorescently tagged mVPS4A mutant localized at the intercellular bridge in both WT and VPS4A KO cells, indicating that VPS4A can be targeted to the intracellular bridge in its monomeric form (Fig 3A). Notably, expression of mVPS4A in VPS4A KO cells partially rescued the abscission phenotype, resulting in a milder delay in abscission and a complete recovery in the percentages of cells that completed abscission (72% in VPS4A KO vs. 91% in mVPS4A OE and 95% in WT cells) (Fig. 3B). Summing the rescue levels measured upon overexpression of VPS4B and mVPS4A resulted an effect that is equivalent to a complete rescue, suggesting that the unique role of VPS4A in abscission, which VPS4B cannot compensate, is carried out by the monomeric form of VPS4A (Fig. 3C).

**Figure 3:**
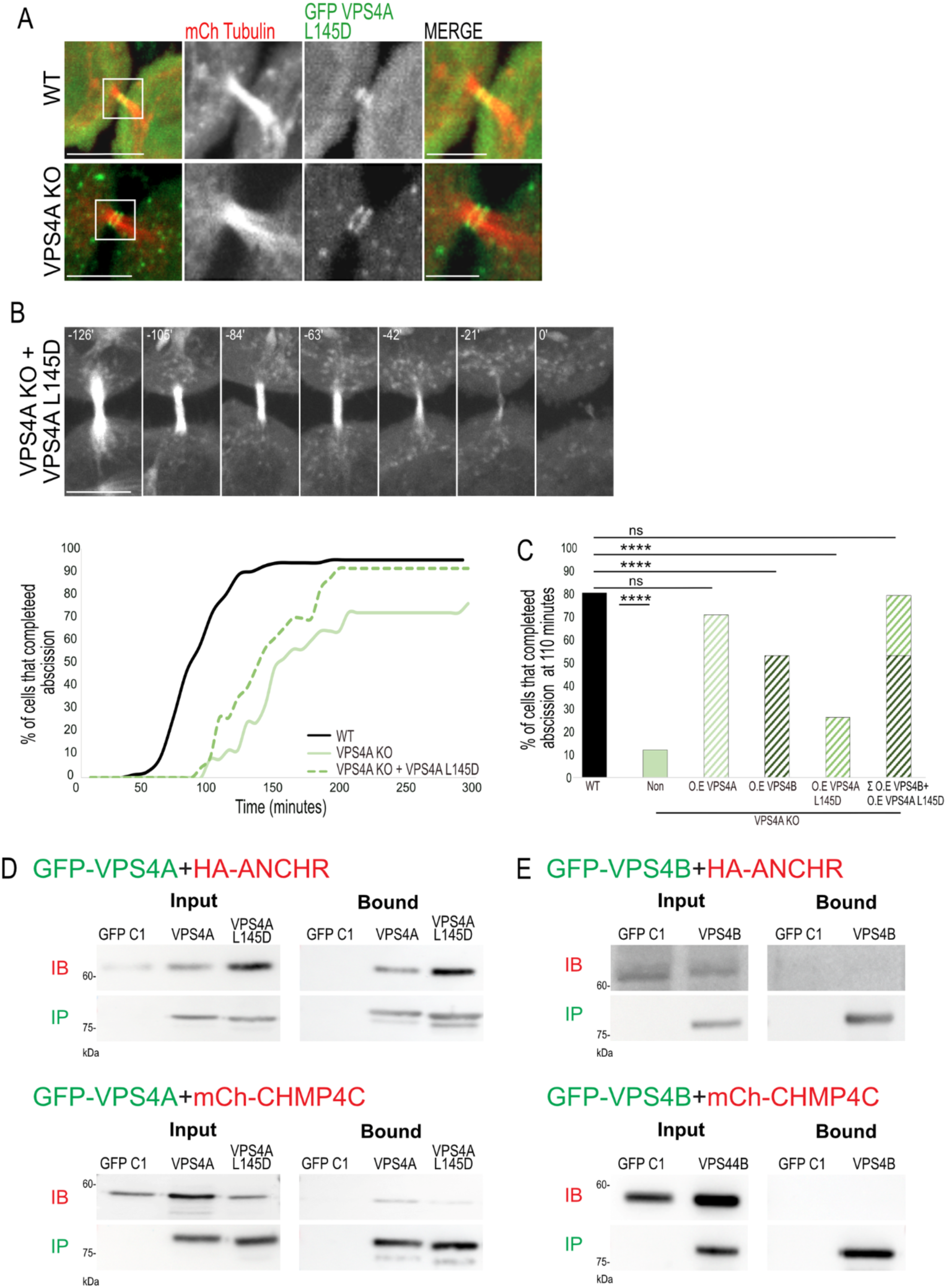
Monomeric VPS4A arrives at the intercellular bridge, partly rescues the abscission delay in VPS4A KO cells, and binds abscission checkpoint proteins. (A) mVPS4A localizes to the intracellular bridge in WT (top panel) and VPS4A KO (bottom panel) cells. Shown are intercellular bridges of cells that were co-transfected with mCherry-tubulin (red) and GFP-mVPS4A (green). Left panels: zoom-out (scale, 10 µm) images. Zoom-in images of the white squares in left panels are shown on the right (scale, 5 µm). Shown are maximum-intensity projection images of representative cells. (B) Expression of a mVPS4A mutant partially rescues the abscission delay in VPS4A KO cells. Top panel: maximum-intensity projection time-lapse images of dividing VPS4A KO cells co-transfected with mCherry-tubulin and GFP-mVPS4A (scale, 10 µm). The abscission time, which was defined as the time of the first microtubule bridge cleavage event, was set as time 0. Bottom left: A cumulative plot showing the duration of cytokinesis (from cleavage furrow formation to microtubule bridge cleavage) of all cells recorded is shown on the right (dashed lines) (VPS4A KO + mVPS4A, 138 ± 32 minutes, n = 23). Data of WT and VPS4A KO cells (reproduced from Fig. 1) are shown for reference (**C**) The fraction of cells that completed abscission at 110 minutes post cleavage furrow as measured for the specified conditions. Note that the sum of partial rescue effects obtained by overexpressing VPS4B and mVPS4A in VPS4A KO cells are equivalent to a complete rescue, suggesting that these two components represent the two distinct functions of VPS4A in abscission. Statistical analysis was performed using Chi-square test ****p ≤ 0.0001. (**D-E**) VPS4A and mVPS4A, but not VPS4B, bind abscission checkpoint proteins. WT cells co-transfected with either GFP-VPS4A, GFP-mVPS4A (left panels), or GFP-VPS4B (right panels) together with HA-ANCHR (top panels) or mCherry-CHMP4C (bottom panels) were subjected to immunoprecipitation using anti-GFP beads. Immunoblotting: IB fractions - anti-HA (top panels) or anti-mCherry (bottom panels); IP fractions - anti-GFP. Data were reproduced in at least two independent experiments.

Previous studies indicated a role for VPS4 in the AuroraB abscission checkpoint (*19, 20, 25*). We therefore asked which of the VPS4 isoforms is part of this checkpoint. In IP experiments, VPS4A and mVPS4A, but not VPS4B, bound the abscission checkpoint proteins ANCHR and CHMP4C (Fig. 3D-E). These data indicate that the regulatory role, previously reported for VPS4 in abscission, is mediated by VPS4A and can potentially be executed by its monomeric form. Our data, therefore, support a dual function for VPS4A in abscission: one that is associated with the canonical VPS4 function as a AAA-ATPase and can potentially be complemented by VPS4B, and another that is regulatory and is independent of its canonical hexameric function.

VTA1 is a co-factor of VPS4 that was shown to stabilize its active, hexameric form (*11, 12, 26-28*). To test whether the regulatory role of VPS4 can be induced by destabilizing the VPS4 hexamer, we examined the effect of VTA1 depletion on VPS4 function in dividing cells using a VTA1 KO cell line (*14*) (Fig. S2A). In native polyacrylamide gel electrophoresis, lower molecular weight bands were observed in VTA1 KO cells compared to WT, supporting decreased levels of hexameric VPS4 in VTA1 KO cells (Fig. S2B). Unexpectedly, loss of VTA1 did not result in a delay in abscission but instead led to accelerated abscission (Fig. 4A). Exogenously expressed VTA1 arrived at the intercellular bridge in VTA1 KO cells and recused the abscission phenotype, indicating that the phenotype is a direct outcome of VTA1 loss (Fig. 4A). The cellular levels of VPS4A and VPS4B were not significantly affected in these cells and both proteins properly localized at the intercellular bridge, indicating that VTA1 is dispensable for targeting VPS4 to the intercellular bridge (Fig. S2C-D). Additionally, no changes in the organization and levels of IST1 at the intercellular bridge were observed, suggesting that the ESCRT-III complex is not significantly affected (Fig. S2E and S2F). Therefore, despite the canonical role of VTA1 in stabilizing the active VPS4 hexamer, its depletion did not significantly affect the arrival and organization of ESCRT-III-VPS4 proteins to the intercellular bridge or lead to delayed abscission.

**Figure 4:**
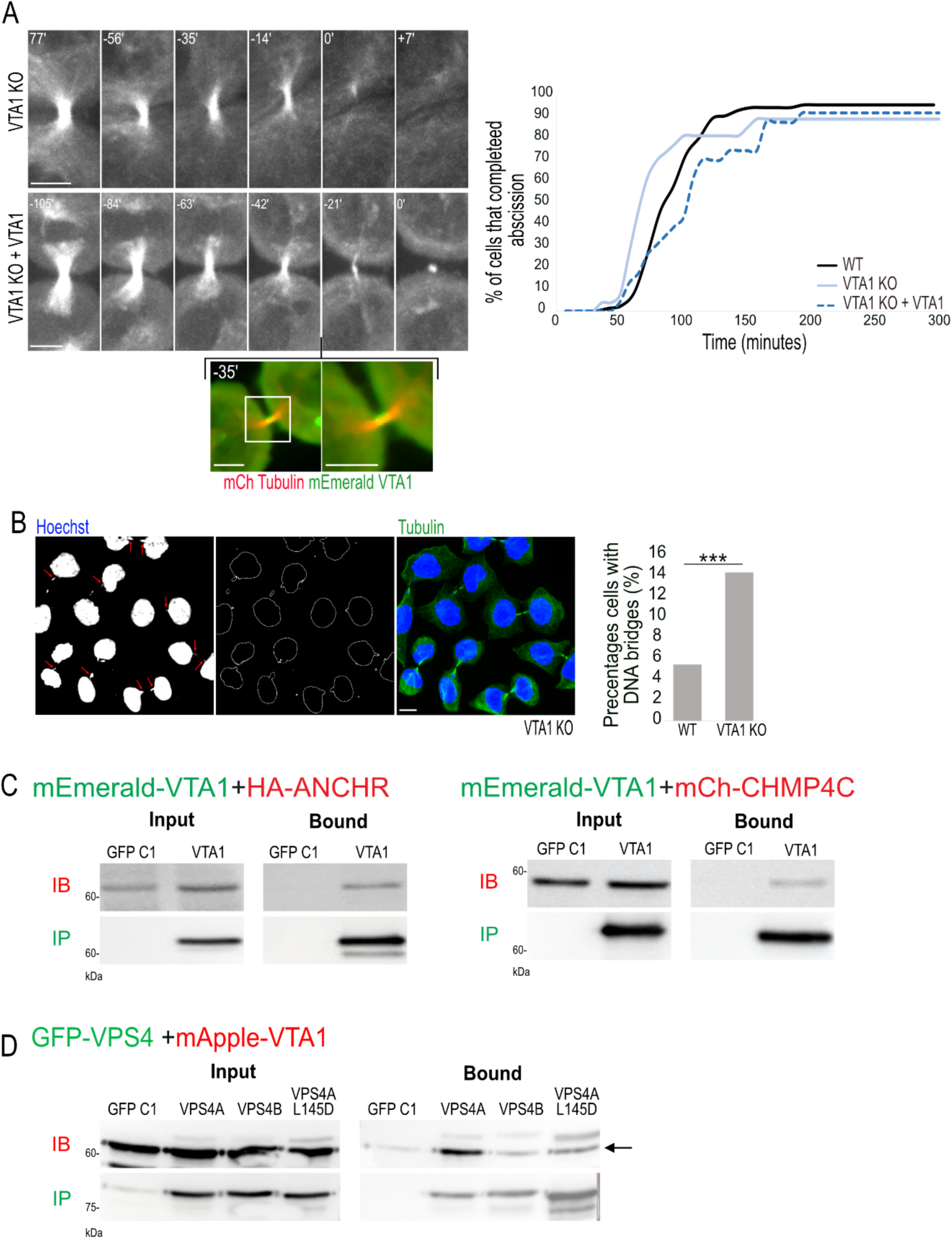
VTA1 is part of the abscission checkpoint complex. (**A**) Abscission is accelerated in cells depleted of VTA1. Live cell imaging of VTA1 KO cells transfected with GFP-tubulin alone (top panel) or with mCherry-tubulin and mEmerald-VTA1 (bottom panel). Shown are maximum-intensity projections of tubulin labeling from time-lapse movies of representative cells. Time 0 = abscission time. Colored images in the inset below demonstrate the arrival of VTA1 to the intercellular bridge during abscission (see also Sup. Movie 3). Scale, 5 µm. A cumulative plot showing the duration of cytokinesis (from cleavage furrow formation to microtubule bridge resolution) of all cells recorded is shown on the right. Averaged duration time of abscission was shorter in VTA1 KO cells compared to WT cells and was recovered by exogenous VTA1 expression (WT 90 ± 46 minutes, n = 76; VTA1 KO 73 ± 28 minutes, n = 26; VTA1 KO + VTA1 102 ± 39 minutes, n = 23). (**B**) Accumulation of DNA bridges in VTA1 KO cells. Representative images of VTA1 KO cells stained with anti-α-tubulin (green) and Hoechst (blue) are shown. Left panel: DNA staining. Arrows indicate DNA bridges. 2^nd^ panel: a Laplacian 2D filter analysis applied to the DNA staining images for better visualization of DNA bridges. 3^rd^ panel: A merged image of tubulin and DNA staining. Scale, 10 µm. Right panel: A plot showing the percentages of cells with DNA bridges in WT and VTA1 KO cells (WT; n = 332, VTA1 KO; n = 287). ***p < 0.001 using chi-square test. (**C**) VTA1 binds the abscission checkpoint proteins CHMP4C and ANCHR. WT cells co-transfected with mEmerald-VTA1 and HA-ANCHR (left panel) or mCherry-CHMP4C (right panel) were subjected to immunoprecipitation (IP) using anti-GFP beads. Immunoblotting: IB fractions anti-HA (left panels) or anti-mCherry (right panels), IP fractions anti-GFP antibodies. (**D**) WT cells co-transfected with the indicated GFP-tagged VPS4 constructs, and mApple-VTA1 were subjected to immunoprecipitation using anti-GFP beads. Immunoblotting: IB fractions anti-VTA1 (Arrow indicates the VTA1 band), IP fractions anti-GFP. Note that VTA1 preferentially binds to VPS4A in cells. Data in C and D were reproduced in at least two independent experiments.

Accelerated abscission was previously reported upon depletion of abscission checkpoint proteins, strongly pointing to the involvement of VTA1 in the abscission checkpoint. Indeed, the level of DNA bridges, a hallmark of checkpoint failure, was significantly increased (Fig. 4B). Levels of the abscission checkpoint protein AuroraB in the intercellular bridge showed no significant difference, suggesting that VTA1 functions downstream to AuroraB (Fig. S2G). In WT cells, VTA1 was found to interact with the checkpoint proteins ANCHR and CHMP4C (Fig 4C). Furthermore, VTA1 could bind both VPS4B and VPS4A, with stronger binding affinity observed for the latter, and was associated with the mVPS4A (Fig 4D). Interestingly, in VTA1 KO cells, VPS4A still exhibited binding to CHMP4C but not to ANCHR, whereas in VPS4A KO cells, VTA1 could bind to ANCHR but not CHMP4C (Fig. 5A-C). These findings strongly indicate that VTA1 is an integral component of the abscission checkpoint and part of CHMP4C/ANCHR/VPS4A complex (Fig. 5D).

**Figure 5:**
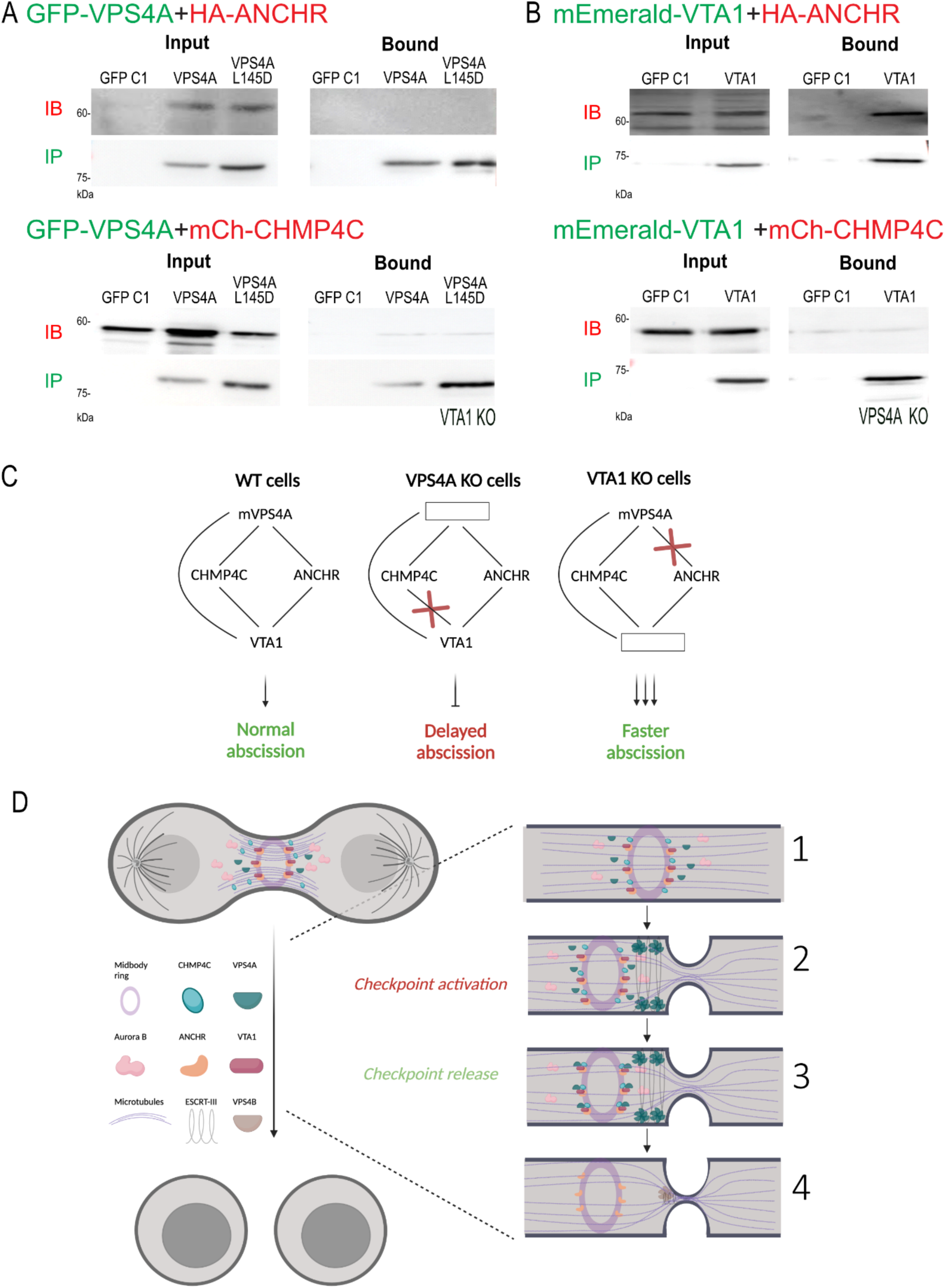
The VPS4A-VTA1 interplay at the abscission checkpoint. (**A-B**) VTA1 KO cells (**A**) or VPS4A KO cells (**B**) co-transfected with the indicated plasmids were subjected to immunoprecipitation (IP) using anti-GFP beads. Immunoblotting: IB fractions anti-HA (top panels) or anti-mCherry (bottom panels), IP fractions anti-GFP. Note that VPS4A-ANCHR interactions were abolished in VTA1 KO cells and VTA1- CHMP4C interactions were abolished in VPS4A KO cells. (**C**) A summary of the interactions between VPS4A and VTA1 and the resulting abscission phenotypes in each of cell lines studied. (**D**) A suggested model for the interplay between VPS4A, VTA1, CHMP4C, and ANCHR at the abscission checkpoint. (1) CHMP4C, ANCHR and VTA1 assemble on the midbody ring with ANCHR and VTA1 forming a complex. (2) VPS4A arrives at the intercellular bridge and assembles into hexamers on the membrane, leading to initial bridge constriction. (3) VPS4A continues accumulating at the bridge, resulting in excess of monomeric VPS4A, which binds to CHMPC at the midbody ring. The latter facilitates the formation of an ANCHR-VTA1-VPS4A-CHMP4C complex at the midbody ring. (5) As a result, the abscission checkpoint is released and ESCRT-driven abscission (mediated by VPS4B) progresses, leading to the completion of abscission and the formation of two independent daughter cells.

## DISCUSSION

In this work, we present the first evidence of a distinct role for VPS4 isoforms in abscission. Our findings support a model in which VPS4B primarily serves the canonical VPS4 function that involves depolymerization of the ESCRT-III filament. At the same time, the VPS4A isoform performs both a canonical role involving ESCRT-III monomer exchange, and the regulatory functions attributed to VPS4 in abscission. We further show that the VPS4A co-factor, VTA1, participates in the VPS4A abscission regulatory checkpoint via binding to checkpoint proteins and facilitating VPS4A-ANCHR interactions. Based on our findings, we propose that VPS4A, which is present in cells at lower levels, acts as a sensor for monitoring the recruitment of VPS4 to the intercellular bridge. When the levels of monomeric VPS4A at the bridge reach a certain threshold, the abscission checkpoint is released, resulting in the completion of abscission. Collectively, our data support a scenario wherein VPS4A and VTA1 cooperate with abscission checkpoint proteins to regulate the timing of abscission (Fig. 5D). Initially, checkpoint proteins CHMP4C, ANCHR, and VTA1 assemble on the midbody ring, with VTA1 and ANCHR forming a complex. VPS4A arrives at the intercellular bridge, assemble into hexamers on the bridge membrane, and drives the initial constriction of the bridge via remodeling the ESCRT-III polymer. As more VPS4A molecules arrive, monomeric VPS4A accumulates at the intercellular bridge and binds to CHMP4C at the midbody ring. This accumulation drives the formation of an ANCHR-VTA1-VPS4A-CHMP4C complex at the midbody ring, which facilitate checkpoint release, possibly by promoting the dephosphorylation of CHMP4C (*25*). Release of the checkpoint allows for the progression and completion of ESCRT-mediated intercellular bridge abscission, forming two fully separated daughter cells. Using the monomeric levels of VPS4A as a readout for VPS4 levels at the intercellular bridge is an elegant mechanism to ensure proper abscission, highlighting the tight regulation of abscission in cells. It will be interesting to examine, in future studies, whether other VPS4-related cellular processes are employing this mode of operation.

Biochemical and structural studies showed that VTA1 stabilizes the VPS4 hexamer and increases ATP hydrolysis by the enzyme (*12, 28*). Our data show that in the absence of VTA1, the arrival of both VPS4 isoforms and ESCRT-III proteins to the intercellular bridge is unperturbed and that abscission is not inhibited. These data suggest that VTA1’s function in stabilizing the VPS4 hexamer is expansible for the function of VPS4 in abscission. On the other hand, our results highlight a fundamental role for VTA1 in the abscission checkpoint. VTA1 was found to bind the checkpoint proteins ANCHR and CHMP4C and to mediate the binding of VPS4A to ANCHR. Furthermore, loss of VTA1 resulted accelerated abscission and accumulation of DNA bridges, two hallmarks of abscission checkpoint failure (*1, 25*). The findings that VTA1 may be expansible for the canonical function of VPS4 in abscission but is a critical factor in the regulation of abscission calls for reinvestigation of the function of VTA1 in cells.

The role of VPS4 in the ESCRT pathway has been studied both in vitro and in cells. Perturbation of VPS4 function inhibited ESCRT-mediated processes in cells, including viral budding and cytokinetic abscission (*3, 14–16, 21, 22, 29, 30*). In vitro, VPS4 was induced the disassembly of ESCRT-III filaments and to drive ESCRT-mediated membrane fission in artificial membrane vesicles, indicating that it is executing the final membrane fission event by depolymerization ESCRT-III filaments (*16, 31-36*). VPS4 also facilitated the exchange of ESCRT-III components in the ESCRT filament, both in vitro and in cells, suggesting an earlier function for VPS4 in shaping the ESCRT-III filament (*16, 36*). Additionally, a role for VPS4 in initiating ESCRT-III assembly was suggested in the context of viral budding and MVB biogenesis (*37, 38*). Finally, VPS4 was found to be involved in the AuroraB checkpoint, which controls the onset of ESCRT mediated cytokinetic abscission (*20*), and to be involved in centrosome maintenance (*4, 5*). How a single protein can drive such diverse functions was unclear. Our STORM analysis suggests temporally separated functions for VPS4A and VPS4B in abscission. While VPS4A is mainly involved in early abscission stages, presumably in monomeric exchange of the ESCRT-III filament, VPS4B appears to have a role later in the process, when ESCRT-III depolymerization occurs. Such functional discrimination between the two VPS4 paralogs provides a potential explanation for the many functions described for VPS4 in cells and raises the possibility that specific VPS4 functions are subjected to different cellular regulations, as demonstrated here for VPS4A. Differences in VPS4A and VPS4B function in cells have been previously reported in the context of viral budding. In particular, it was shown that the HIV virus mainly utilizes the VPS4B isoform to bud out of cells (*14, 15*). Our observations that the cellular levels of VPS4B are significantly higher than VPS4A and that VPS4A is involved in the regulation of abscission may provide a logical explanation for these observations. Utilizing the VPS4B isoform is advantageous for the virus, as this isoform is more abundant in the cells and less involved in cellular regulation.

VPS4 has been previously associated with various pathologies, including cancer and neurodevelopmental disorders. However, while the deletion of VPS4B has been linked to cancer mostly indirectly due to its location on chromosome 18, which contains crucial tumor suppressor genes (*8*), specific heterozygous point mutations in VPS4A alone were sufficient to induce severe pathologies (*6, 7*). Two independent studies have demonstrated that individuals carrying single mutations in the VPS4A gene developed Dyserythropoietic Anemia and severe neurodevelopmental delays (*6, 7*). These conditions were associated with structural brain abnormalities, intellectual disability (ID), deafness, cataracts, and visual dysfunction. In one of the studies, fibroblasts taken from patients showed defects in centrosome number, primary cilia morphology, and cell cycle progression (*6*). Interestingly, EGFR degradation, which relies on ESCRTs for internalization to multivesicular bodies, remained normal, suggesting that specific ESCRT-dependent cellular processes are more sensitive to the loss of VPS4A. Notably, enlarged endosomes with abnormal IST1 accumulations, but not other ESCRT-III proteins, were observed in these cells, strongly indicating a link between VPS4A and IST1. These findings are consistent with our STORM data, which demonstrated that IST1 accumulates at the intercellular bridge upon VPS4A loss. In the second study, blood cells from patients displayed bi-nucleated cells and accumulation of cytokinetic bridges (*7*). iPSC cells derived from these patients exhibited late cytokinesis defects accompanied by mislocalization of VPS4 at the intercellular bridge. The crucial role of VPS4A in neurodevelopment suggests that its cellular regulation is paramount for proper neuronal development and function. Despite the presence of a normal VPS4B gene and adequate VPS4B levels in these patients, the disruption of VPS4A seems to impact disease progression significantly, indicating the uniqueness of VPS4A’s functions in these pathologies. However, so far, a mechanistic explanation for these observations has been lacking. Our findings provide, for the first time, a potential mechanistic explanation for these observations. We propose that patients who carry normal VPS4B and mutated VPS4A genes develop pathologies due to the disruption of VPS4A-mediated cellular regulation, which is essential for neurodevelopment and cannot be compensated by VPSB. This regulation of VPS4A may be related to the abscission regulation reported here or other regulatory roles of VPS4A that are yet to be characterized. Further investigation of the regulatory functions of VPS4A in cells could open new avenues for targeted therapeutic interventions for these severe pathologies.

## METHODS

### Cell culture and transfection

HeLa cells were grown in DMEM supplemented DMEM with 10% fetal bovine serum (FBS), 2mM glutamine, 10,000 U/mL penicillin, 10 mg/mL streptomycin at 37°C and 5% CO2. Transfection was carried out by using Lipofectamine 2000 (Life Technologies), PolyJet (SignaGen Laboratories) or JetPrime transfection kit (Polyplus), according to manufacturer’s guidelines.

### Plasmid constructs

Fluorescently tagged versions of human VPS4A and VPS4B were previously cloned in our laboratory as described in Ott et. al., (*5*). A monomeric VPS4A version (L145D) was designed based on the previously published monomeric yeast VPS4 (VPS4 L151D) (*24*). The point mutation at position L145 was introduced by overlapping PCR. GFP-ANCHR: Full-length human ANCHR conjugated to GFP was a kind gift by Harald Stenmark (Centre for Cancer Biomedicine, Faculty of Medicine, Oslo University Hospital, Norway) (*20*). An HA tagged version (HA-ANCHR) was generated by subcloning ANCHR sequence into an 3xHA-C1 plasmid. mEmerald/ mApple -VTA1: full-length human VTA1 was amplified by PCR and cloned to mEmerald or mApple -C1 vectors (Clontech). mCh-CHMP4C, GFP–α-tubulin, and mCherry–α-tubulin were generated as previously described (*18*).

All constructs were confirmed by sequencing.

### Genome editing

For CRISPR/Cas9-mediated gene disruption, guide RNAs (gRNAs) Guide RNAs (gRNAs) targeting VPS4A, VPS4B, and VTA1 were sub-cloned into the lentiCRISPR plasmid (Addgene, #49535). Following transfection and puromycin selection, single clones were isolated and expanded. To confirm the efficacy of protein knockouts, Western Blot analysis was performed. The gRNA sequences employed were as follows: CACCGAGTGCGTGCAGTACCTAGAC for targeting VPS4A, CACCGCAAACAGAAAGCGATAGATC for targeting VPS4B, and CACCGGCATGACAAGCGAGACCCTG for targeting VTA1.

### Cell Synchronization

Double thymidine block was performed as previously described (*21*). In short, HeLa cells were plated at 10% confluency on an Ibidi Glass Bottom Dish 35 mm (Martinsried, Germany) and treated with 2 mM thymidine (Sigma, T1895-1G) for 18 hours to induce the first block. After incubation, cells were washed once with PBS and grown in fresh media for 9 hours. Cells were then supplemented for an additional 15 hours with 2 mM thymidine, washed once with PBS, and grown in fresh media for 10.5-12.5 hours to enrich the percentages of intercellular bridges.

### Immunoprecipitation (IP)

Immunoprecipitation was carried out using the GFP-Trap Agarose KIT (ChromoTek), following the manufacturer’s instructions with minor modifications. Transfected HeLa cells were harvested 24 hours after transfection and washed with PBS. The cells were then lysed on ice for 30 minutes using a lysis buffer containing 10 mM Tris/Cl (pH 7.5), 100 mM NaCl, 0.5 mM EDTA, 0.5% NP40, 1 mM PMSF, and a complete protease inhibitor cocktail (Roche Diagnostics). Cell lysates were then centrifuged at 20,000 g for 10 minutes at 4°C, and supernatants were diluted in a buffer consisting of 10 mM Tris/Cl (pH 7.5), 150 mM NaCl, 0.5 mM EDTA, 1 mM PMSF, and a complete protease inhibitor cocktail. The diluted lysates were incubated with GFP-Trap-A beads for 1 hour. Subsequently, the lysates and beads were centrifuged at 2,500 g for 2 minutes at 4°C to separate the unbound fraction from the bound fraction. Samples were heated at 95°C for 5 min in Laemmli sample buffer and resolved by SDS-PAGE followed by western blot analysis.

### Western blot

HeLa cells were lysed as described above or in RIPA lysis buffer [150 mM NaCl, 1% NP-40, 0.5% deoxycholate, 0.1% SDS, 50 mM Tris (pH 8.0)] supplemented with complete protease inhibitor (Roche Diagnostics) for 30 min at 4 °C. Total protein concentrations of the lysates were determined using BCA Protein Assay Kit (Pierce Biotechnology). Samples were heated at 95°C for 5 min in Laemmli sample buffer and resolved by SDS-PAGE. Membranes were stained with primary antibodies for 16 hours at 4 °C using the following antibodies: anti-GFP (Applied Biological Materials, G096, 1:1000), anti mCherry (novus biologicals, NBP2-25157, 1:1000-WB), anti VTA1 (ThermoFisher, PA5-21831 1:1500), anti-HA (Applied Biological Materials, G166,1:4000), anti-VPS4 (Sigma-Aldrich, SAB4200025, 1:500). Then a secondary anti-rabbit or anti-mouse-peroxidase antibodies (1:10,000; Jackson ImmunoResearch) were applied for 1 hour.

### Native PAGE

HeLa wild-type (WT) and VTA1 knockout (KO) cells were lysed using a hypotonic buffer consisting of 10 mM HEPES (pH 7.9), 1.5 mM MgCl, and 10 mM KCl. A total of 30 μl cell extracts were loaded on native polyacryamide gel. The membrane was then incubated with primary antibodies overnight at 4°C, followed by incubation with anti-rabbit peroxidase-conjugated secondary antibodies for 1 hour at 1:10,000 dilution (Jackson ImmunoResearch).

### Immunofluorescence

HeLa cells were washed with PBS, fixed with 4% paraformaldehyde (PFA), permeabilized with 0.5% Triton X-100 for 10 min, and blocked with 10% FBS for 15 min. Fixed cells were stained with the primary antibodies: anti-CHMP4B (Proteintech, 13683- 1-AP, 1:50), anti-IST1 (Proteintech, 51002-1-AP, 1:50), anti-tubulin (Sigma, T6199 1:1000), anti-AurB (abcam, ab2254, 1:1000) and anti-AurBpT232 (Rockland, 600-401- 667S, 1:500-IF), as indicated, and subjected to fluorescently-tag secondary antibodies staining (Alexa Fluor 488 or Alexa Fluor 594, Life Technologies). Finally, cells were mounted with Fluoromount-G (SouthernBiotech, Birmingham, AL) and imaged.

### Live Cell Imaging

HeLa cells were plated at low density in four-well chamber slides (Ibidi, Martinsried, Germany), transfected 24 hours later with the indicated plasmids, and imaged 24–48 hours later. Z-stacks of selected low-expressing cells undergoing cytokinesis were collected at the specified intervals using a fully incubated confocal spinning-disk microscope (Marianas; Intelligent Imaging, Denver, CO) with a 63× oil objective (numerical aperture, 1.4) and were video recorded on an electron-multiplying charge-coupled device camera. (Evolve; (pixel size, 0.079 μm; Evolve; Photometrics). Image processing and analysis were done using SlideBook version 6 (3I Inc).

### Structured Illumination Microscopy (SIM)

Cells were plated at low density on high-resolution #1.5 coverslips (Marienfeld, Lauda-Konigshfen, Germany) and fixed using 4% PFA. Cells were further subjected to immunostaining as described above. 3D SIM imaging of low expressing cells was performed using the ELYRA PS.1 microscope (Carl Zeiss MicroImaging). Thin z-sections (0.11–0.15 μM) of high-resolution images were collected in three rotations and five phases for each channel. Image reconstruction and processing were performed in ZEN (Carl Zeiss MicroImaging).

### STORM analysis

Synchronized cells were washed with PBS, fixed with 3% paraformaldehyde (PFA) and 0.1% glutaraldehyde, and permeabilized and blocked with 0.2% Triton X-100 with 3% BSA. Cells were then stained with tubulin and IST1 antibodies. Anti-mouse Alexa Fluor 488 and anti-rabbit Alexa Fluor 647 (Life Technologies, 1:1500) were used as secondary antibodies, respectively. Cells were subjected to a second fixation (as above). STORM acquisition was performed in STORM buffer containing 7 μM glucose oxidase (Sigma), 56 nM catalase (Sigma), 5 mM cysteamine (Sigma), 50 mM Tris, 10 mM NaCl, and 10% glucose (pH 8) using a Zeiss Elyra ELYRA PS.1 microscope in wide-field mode (100X N.A 1.46 oil immersion objective. Excitation was induced using a 647 nm laser (5 kW/cm^2^). A total of 10,000 frames were acquired for each dataset with an acquisition time of 35 ms per frame. STORM analysis was done using ZEN 2011 software (Zeiss, Germany). Datasets were filtered similarly for all conditions according to the following parameters: peak mask size – 5, peak intensity to noise – 7, drift correction – 3, localization precision – 1-30nm, minimal number of photons – 500, first frame – 1000 and up. Localizations densities at the intercellular bridge were calculated as the ratio between the number of localizations and the area of an ROI surrounding the structure which was depicted manually (# of localizations/area).

### Statistical analysis

Statistical analysis was performed using Graph Pad Prism version 9.00 for Windows (La Jolla, CA, USA). Normality was tested using the D’Agostino-Pearson test. For non-normal distributions, statistical significance between two groups was determined using the unpaired two-tailed Mann-Whitney U test and between multiple groups using the unpaired one-way analysis of variance (ANOVA) Dunn’s test. For normal distributions, statistical significance between two groups was determined using two-tailed T-test or by chi-square test, and between the multiple groups was determined using the paired one-way analysis of variance (ANOVA) tukey’s test. Statistical Significance in all plots is as follows: ns (non-significant) p-value> 0.05, *p-value ≤ 0.05, **p-value ≤ 0.01, ***p-value ≤ 0.001, ****p-value ≤ 0.0001. Bars in all plots indicate Standard deviation of the dataset.

Illustrations were generated in Biorender.

## ACKNOWLEDGMENTS

We thank all members of the Elia lab for critical feedback throughout the project. The Elia laboratory is funded by the Israeli Science Foundation (ISF) Grant no. 1323/18.

## AUTHOR CONTRIBUTIONS

DN and NE conceptualized the project. ID performed and analyzed all experiments using VTA1 KO cells. YA performed and analyzed all experiments using VPS4 KO cells. ID and DN performed and analyzed all IP and biochemical assays. DN and YA generated KO cell lines. ID prepared figures for publication. NE wrote the manuscript with the help of DN. All authors read and revised the manuscript.

Competing Interest Statement: We declare no competing interests.

## REFERENCES

1. A. T. Gatta, J. G. Carlton, The ESCRT-machinery: closing holes and expanding roles. Curr Opin Cell Biol 59, 121–132 (2019).

2. J. H. Hurley, ESCRTs are everywhere. EMBO J 34, 2398–2407 (2015).

3. Y. A. M. Alonso, S. M. Migliano, D. Teis, ESCRT-III and Vps4: a dynamic multipurpose tool for membrane budding and scission. FEBS J 283, 3288–3302 (2016).

4. E. Morita, L. A. Colf, M. A. Karren, V. Sandrin, C. K. Rodesch, W. I. Sundquist, Human ESCRT-III and VPS4 proteins are required for centrosome and spindle maintenance. Proc Natl Acad Sci U S A 107, 12889–12894 (2010).

5. C. Ott, D. Nachmias, S. Adar, M. Jarnik, S. Sherman, R. Y. Birnbaum, J. Lippincott-Schwartz, N. Elia, VPS4 is a dynamic component of the centrosome that regulates centrosome localization of gamma-tubulin, centriolar satellite stability and ciliogenesis. Sci Rep 8, 3353 (2018).

6. C. Rodger, E. Flex, R. J. Allison, A. Sanchis-Juan, M. A. Hasenahuer, S. Cecchetti, C. E. French, J. R. Edgar, G. Carpentieri, A. Ciolfi, F. Pantaleoni, A. Bruselles, C. Genomics England Research, R. Onesimo, G. Zampino, F. Marcon, E. Siniscalchi, M. Lees, D. Krishnakumar, E. McCann, D. Yosifova, J. Jarvis, M. C. Kruer, W. Marks, J. Campbell, L. E. Allen, S. Gustincich, F. L. Raymond, M. Tartaglia, E. Reid, De Novo VPS4A Mutations Cause Multisystem Disease with Abnormal Neurodevelopment. Am J Hum Genet 107, 1129–1148 (2020).

7. K. G. Seu, L. R. Trump, S. Emberesh, R. B. Lorsbach, C. Johnson, J. Meznarich, H. R. Underhill, S. T. Chou, H. Sakthivel, N. N. Nassar, K. J. Seu, L. Blanc, W. Zhang, C. M. Lutzko, T. A. Kalfa, VPS4A Mutations in Humans Cause Syndromic Congenital Dyserythropoietic Anemia due to Cytokinesis and Trafficking Defects. Am J Hum Genet 107, 1149–1156 (2020).

8. E. Szymanska, P. Nowak, K. Kolmus, M. Cybulska, K. Goryca, E. Derezinska-Wolek, A. Szumera-Cieckiewicz, M. Brewinska-Olchowik, A. Grochowska, K. Piwocka, M. Prochorec-Sobieszek, M. Mikula, M. Miaczynska, Synthetic lethality between VPS4A and VPS4B triggers an inflammatory response in colorectal cancer. EMBO Mol Med 12, e10812 (2020).

9. J. X. Wei, L. H. Lv, Y. L. Wan, Y. Cao, G. L. Li, H. M. Lin, R. Zhou, C. Z. Shang, J. Cao, H. He, Q. F. Han, P. Q. Liu, G. Zhou, J. Min, Vps4A functions as a tumor suppressor by regulating the secretion and uptake of exosomal microRNAs in human hepatoma cells. Hepatology 61, 1284–1294 (2015).

10. H. Han, J. M. Fulcher, V. P. Dandey, J. H. Iwasa, W. I. Sundquist, M. S. Kay, P. S. Shen, C. P. Hill, Structure of Vps4 with circular peptides and implications for translocation of two polypeptide chains by AAA+ ATPases. Elife 8, (2019).

11. S. Sun, L. Li, F. Yang, X. Wang, F. Fan, M. Yang, C. Chen, X. Li, H. W. Wang, S. F. Sui, Cryo-EM structures of the ATP-bound Vps4(E233Q) hexamer and its complex with Vta1 at near-atomic resolution. Nat Commun 8, 16064 (2017).

12. I. F. Azmi, B. A. Davies, J. Xiao, M. Babst, Z. Xu, D. J. Katzmann, ESCRT-III family members stimulate Vps4 ATPase activity directly or via Vta1. Dev Cell 14, 50–61 (2008).

13. S. Scheuring, R. A. Rohricht, B. Schoning-Burkhardt, A. Beyer, S. Muller, H. F. Abts, K. Kohrer, Mammalian cells express two VPS4 proteins both of which are involved in intracellular protein trafficking. J Mol Biol 312, 469–480 (2001).

14. S. Harel, Y. Altaras, D. Nachmias, N. Rotem-Dai, I. Dvilansky, N. Elia, I. Rousso, Analysis of individual HIV-1 budding event using fast AFM reveals a multiplexed role for VPS4. Biophys J 121, 4229–4238 (2022).

15. C. Kieffer, J. J. Skalicky, E. Morita, I. De Domenico, D. M. Ward, J. Kaplan, W. I. Sundquist, Two distinct modes of ESCRT-III recognition are required for VPS4 functions in lysosomal protein targeting and HIV-1 budding. Dev Cell 15, 62–73 (2008).

16. B. E. Mierzwa, N. Chiaruttini, L. Redondo-Morata, J. M. von Filseck, J. Konig, J. Larios, I. Poser, T. Muller-Reichert, S. Scheuring, A. Roux, D. W. Gerlich, Dynamic subunit turnover in ESCRT-III assemblies is regulated by Vps4 to mediate membrane remodelling during cytokinesis. Nat Cell Biol 19, 787–798 (2017).

17. N. Elia, C. Ott, J. Lippincott-Schwartz, Incisive imaging and computation for cellular mysteries: lessons from abscission. Cell 155, 1220–1231 (2013).

18. N. Elia, R. Sougrat, T. A. Spurlin, J. H. Hurley, J. Lippincott-Schwartz, Dynamics of endosomal sorting complex required for transport (ESCRT) machinery during cytokinesis and its role in abscission. Proc Natl Acad Sci U S A 108, 4846–4851 (2011).

19. V. Nahse, L. Christ, H. Stenmark, C. Campsteijn, The Abscission Checkpoint: Making It to the Final Cut. Trends Cell Biol 27, 1–11 (2017).

20. S. B. Thoresen, C. Campsteijn, M. Vietri, K. O. Schink, K. Liestol, J. S. Andersen, C. Raiborg, H. Stenmark, ANCHR mediates Aurora-B-dependent abscission checkpoint control through retention of VPS4. Nat Cell Biol 16, 550–560 (2014).

21. I. Goliand, S. Adar-Levor, I. Segal, D. Nachmias, T. Dadosh, M. M. Kozlov, N. Elia, Resolving ESCRT-III Spirals at the Intercellular Bridge of Dividing Cells Using 3D STORM. Cell Rep 24, 1756–1764 (2018).

22. J. Guizetti, L. Schermelleh, J. Mantler, S. Maar, I. Poser, H. Leonhardt, T. Muller-Reichert, D. W. Gerlich, Cortical constriction during abscission involves helices of ESCRT-III-dependent filaments. Science 331, 1616–1620 (2011).

23. N. Monroe, H. Han, M. D. Gonciarz, D. M. Eckert, M. A. Karren, F. G. Whitby, W. I. Sundquist, C. P. Hill, The oligomeric state of the active Vps4 AAA ATPase. J Mol Biol 426, 510–525 (2014).

24. M. D. Gonciarz, F. G. Whitby, D. M. Eckert, C. Kieffer, A. Heroux, W. I. Sundquist, C. P. Hill, Biochemical and structural studies of yeast Vps4 oligomerization. J Mol Biol 384, 878–895 (2008).

25. J. G. Carlton, A. Caballe, M. Agromayor, M. Kloc, J. Martin-Serrano, ESCRT-III governs the Aurora B-mediated abscission checkpoint through CHMP4C. Science 336, 220–225 (2012).

26. N. Monroe, H. Han, P. S. Shen, W. I. Sundquist, C. P. Hill, Structural basis of protein translocation by the Vps4-Vta1 AAA ATPase. Elife 6, (2017).

27. D. Yang, J. H. Hurley, Structural role of the Vps4-Vta1 interface in ESCRT-III recycling. Structure 18, 976–984 (2010).

28. I. Azmi, B. Davies, C. Dimaano, J. Payne, D. Eckert, M. Babst, D. J. Katzmann, Recycling of ESCRTs by the AAA-ATPase Vps4 is regulated by a conserved VSL region in Vta1. J Cell Biol 172, 705–717 (2006).

29. C. Caillat, S. Maity, N. Miguet, W. H. Roos, W. Weissenhorn, The role of VPS4 in ESCRT-III polymer remodeling. Biochem Soc Trans 47, 441–448 (2019).

30. G. H. Pieper, S. Sprenger, D. Teis, S. Oliferenko, ESCRT-III/Vps4 Controls Heterochromatin-Nuclear Envelope Attachments. Dev Cell 53, 27–41 e26 (2020).

31. B. Yang, G. Stjepanovic, Q. Shen, A. Martin, J. H. Hurley, Vps4 disassembles an ESCRT-III filament by global unfolding and processive translocation. Nat Struct Mol Biol 22, 492–498 (2015).

32. J. Schoneberg, M. R. Pavlin, S. Yan, M. Righini, I. H. Lee, L. A. Carlson, A. H. Bahrami, D. H. Goldman, X. Ren, G. Hummer, C. Bustamante, J. H. Hurley, ATP-dependent force generation and membrane scission by ESCRT-III and Vps4. Science 362, 1423–1428 (2018).

33. S. Maity, C. Caillat, N. Miguet, G. Sulbaran, G. Effantin, G. Schoehn, W. H. Roos, W. Weissenhorn, VPS4 triggers constriction and cleavage of ESCRT-III helical filaments. Sci Adv 5, eaau7198 (2019).

34. S. Lata, G. Schoehn, A. Jain, R. Pires, J. Piehler, H. G. Gottlinger, W. Weissenhorn, Helical structures of ESCRT-III are disassembled by VPS4. Science 321, 1354–1357 (2008).

35. N. de Franceschi, M. Alqabandi, W. Weissenhorn, P. Bassereau, Dynamic and Sequential Protein Reconstitution on Negatively Curved Membranes by Giant Vesicles Fusion. Bio Protoc 9, e3294 (2019).

36. A. K. Pfitzner, V. Mercier, X. Jiang, J. Moser von Filseck, B. Baum, A. Saric, A. Roux, An ESCRT-III Polymerization Sequence Drives Membrane Deformation and Fission. Cell 182, 1140–1155 e1118 (2020).

37. M. A. Y. Adell, S. M. Migliano, S. Upadhyayula, Y. S. Bykov, S. Sprenger, M. Pakdel, G. F. Vogel, G. Jih, W. Skillern, R. Behrouzi, M. Babst, O. Schmidt, M. W. Hess, J. A. Briggs, T. Kirchhausen, D. Teis, Recruitment dynamics of ESCRT-III and Vps4 to endosomes and implications for reverse membrane budding. Elife 6, (2017).

38. D. S. Johnson, M. Bleck, S. M. Simon, Timing of ESCRT-III protein recruitment and membrane scission during HIV-1 assembly. Elife 7, (2018).

